# Non-linear effects of socioeconomic status on brain development: associations between parental occupation, cortical thickness and language skills in childhood and adolescence

**DOI:** 10.1101/575993

**Authors:** Budhachandra Khundrakpam, Suparna Choudhury, Uku Vainik, Noor Al-Sharif, Neha Bhutani, Alan Evans, the Pediatric Imaging, Neurocognition and Genetics Study

**Affiliations:** Montreal Neurological Institute, McGill University, 3801 Rue University, Montreal, Quebec, Canada H3A 2B4; Division of Social & Transcultural Psychiatry, McGill University, 1033 Pine Avenue West, Montreal, Quebec, Canada H3A 1A1; Ludmer Centre for Neuroinformatics & Mental Health, McGill University, 3661 Rue University, Montreal, Quebec, Canada H3A 2B3

**Keywords:** Cortical thickness, socioeconomic status (SES), parental occupation, poverty, brain development, adolescence, language, moderated mediation model, PING

## Abstract

Studies have pointed to the role of the brain in mediating the effects of the social environment of the developing child on life outcomes. Since brain development involves nonlinear trajectories, these effects of the child’s social context will likely have age-related differential associations with the brain. However, there is still a dearth of integrative research investigating the interplay between neurodevelopmental trajectories, social milieu and life outcomes. We set out to fill this gap, focusing specifically on the role of socioeconomic status, SES (indexed by parental occupation) on brain and cognitive development by analyzing MRI scans from 757 typically-developing subjects (age = 3-21 years). We observed nonlinear interaction of age and SES on cortical thickness, specifically a significant positive association between SES and thickness around 9-13 years at several cortical regions. Using a moderated mediation model, we observed that cortical thickness mediated the link between SES and language abilities, and this mediation was moderated by ‘age’ in a quadratic pattern, indicating a pronounced SES-effect during early adolescence. Our results, drawn from cross-sectional data, provide a basis for further longitudinal studies to test whether early adolescence may be a sensitive time window for the impact of SES on brain and cognitive development.

## Introduction

The experience of living with socioeconomic disadvantage during childhood and adolescence has been consistently linked to pronounced differences in mental and physical health, educational attainment, cognitive and social-emotional development (Brooks-Gunn and Duncan 1997; Ackerman et al. 2004). Poorer families are more likely to experience low birth weight babies, birth defects, fetal alcohol syndrome among other problems, mediated by processes ranging from the experience of racism to poor maternal nutrition and toxic environments in the neighbourhood (Aber et al. 1997). During childhood, poverty is associated with higher rates of respiratory illnesses, infections (Coultas et al. 1994); while lower cognitive development including academic attainment levels at school (Welsh et al. 2010) as well as socio-emotional adjustment problems have been observed during middle childhood (Brooks-Gunn and Duncan 1997). Studies have also pointed to greater incidence of mental health problems such as depression among adolescents from poorer families (McLoyd 1998; Sander and McCarty 2005; Thapar et al. 2012). These data suggest that the developing brain is shaped by the social ecology in which young people live, and as such, have implications for life outcomes.

A major challenge for researchers studying the health impacts of poverty concerns the conceptual and operational definition of poverty, itself, and in particular, childhood poverty (Minujin et al. 2006). Researchers conducting empirical studies generally use socioeconomic status (SES) as a proxy for poverty. SES is an indicator of a family’s access to social and economic resources, and the advantages and social status these resources allow for (Brito and Noble 2014; Farah 2017). SES, as it is operationalized in quantitative studies, is a multidimensional construct, most commonly estimated by some permutation of three objective components, which, when concerned with SES of children pertain to the parent(s): income, occupation and education level. Subjective measures of social standing and neighborhood quality are often considered in SES measures as well. As such SES not only reflects economic resources but also aspects of social hierarchy and prestige.

A burgeoning literature in developmental cognitive neuroscience has begun to address the question of how SES impacts on neurocognitive development. However, we observe that neuroscience studies, thus far, have focused on correlating structural and functional MRI data with either composite measures of SES, family income or parental education, giving little attention to the component of parental occupation. Parental occupation may be an important and neglected indicator of childhood and adolescent SES compared to absolute measures of material resources or academic attainment because, while related, it may more precisely capture position in the social hierarchy, which has consistently been shown to be intimately related to health and life chances. Most notably, the Whitehall Study, a classic large scale social epidemiological study of British civil servants that begun in 1967, demonstrated that occupational grade is inversely associated with mortality from a range of diseases (Marmot et al. 1978, 1991). Indeed, social stratification is often conceptualized in terms of one’s job, impacting health through accessible privileges (e.g. health care) and social standing in terms of the positional relation to others in the social structure (Galobardes et al. 2007). A parent’s experience of social rank can structure the home environment and parenting style in ways that are transmitted to, and felt by, the child, especially at developmental periods when social status is particularly salient. Indeed, data from the social determinants of health literature have shown that parental occupation has a direct impact on the health and educational attainment of offspring (Pinilla et al. 2017) and that stress during childhood due to parents’ occupational status is linked to risk of later cardiovascular problems in adulthood (Deschênes et al. 2018). We therefore set out to build on findings in the social determinants of health literature and examine the relationship between parental occupation and health outcomes, specifically, neurocognitive development.

Our goal is also to better characterize the relationship between SES, brain structure, cognition and age, an area which has important implications for understanding trajectories of brain-environment interactions and for shaping policy related to social inequality. Given that typical brain development involves intricate processes with regionally specific nonlinear trajectories (Giedd et al. 1999; Gogtay et al. 2004; Shaw et al. 2008) and that SES is a complex, multifactorial phenomenon likely to have differential effects at different time points, it is possible that the brain-SES relation may vary *non-linearly* with age. Put differently, lower SES, for example, may render a person more or less vulnerable to the effects on cognition and mental health at different periods of her development. It is likely that the brain’s trajectory of maturation over age mediates these greater or lesser impacts. An existing study that draws on data from the Pediatric Imaging, Neurocognition and Genetics (PING) study (http://pingstudy.ucsd.edu/Data.php), investigated children and adolescents (age: 5-17 years) and showed *linear* (SES × age) interaction in the left superior temporal gyrus and left inferior frontal gyrus, with a positive relationship between SES and volume emerging in adolescence (Noble et al. 2012). The same group, using a larger sample of participants from the PING dataset (1148 children and adolescents), observed a *curvilinear* association of age and cortical thickness for children from lower SES families while children from higher SES families showed linear association of age and cortical thickness (Piccolo et al. 2016). However, this *non-linear* (SES × age) interaction was observed for the average cortical thickness (of all brain regions) whereas at region-level, this non-linear interaction was not observed (read as, not significant) except at the left fusiform gyrus which the authors showed using post-hoc analysis. Another study by the same group looked at the three-way interactions of age^2^, average cortical thickness and SES (family income) and observed near significance (*p* = 0.07) for executive functions (Brito et al. 2017).

Our study therefore aims to shed new light on the relationship between SES and brain development in two specific ways. Given that neurodevelopmental trajectories show high regional specificity (Gogtay et al. 2004; Shaw et al. 2008) which in turn relate to cognition (Shaw et al. 2006), it is important to further investigate the non-linear (SES × age) interaction on cortical thickness at region-level, and its relation to cognitive development. Additionally, existing studies that draw on the PING dataset have used family income and parental education as measures of SES. Since different SES measures have differential impacts on brain structure and cognition (Noble et al. 2015; Brito et al. 2017), investigating parental occupation as a measure of SES may provide distinct patterns of interaction with brain structure and cognition. We therefore set out to explore *non-linear* (SES × age) interaction (with SES as measured by parental occupation) with cortical thickness. Additionally, using scores on language abilities (vocabulary and reading scores), we set out to test whether the non-linear (SES × age) interaction with cortical thickness relate to differential life outcomes (in terms of language abilities). Lastly, we set out to build a moderated mediation model to integrate all our findings and explain their relationships.

## Materials and Methods

### Subjects

The data for the study were obtained from the Pediatric Imaging, Neurocognition and Genetics (PING) study (http://pingstudy.ucsd.edu/Data.php). The PING study (Jernigan et al. 2016) is a comprehensive, publicly shared, data resource for investigating neurocognition, neuroimaging and genetics in normally developing children and adolescents. The cohort, details described elsewhere (Akshoomoff et al. 2014; Jernigan et al. 2016) comprised of cross-sectional measurements on 1493 subjects (aged 3-21 years) aggregated from sites across the United States. Each subject’s medical, developmental, behavioral history as well as family medical history and environment were obtained from parental questionnaires. Socio-economic status (SES) was recorded as a seven-point scale rating parental occupation from “unskilled employees” to “higher executives”, seven-point scale rating parental education from “less than seven years” to “professional degree”, and a twelve-point scale rating annual familial income from “less than $5,000” to “over $300,000”. Neurocognitive abilities were assessed using the NIH Toolbox Cognition Battery (NTCB, http://www.nihtoolbox.org/) a computerized battery designed for administration across the lifespan (Akshoomoff et al. 2014)). The NTCB includes eight subtests spanning six domains. For our study, we used two language measures (Picture Vocabulary and Oral Reading Recognition tests) that were used in earlier studies (Brito et al. 2017; Schork et al. 2018).

### Image acquisition and pre-processing

Each site administered a standardized structural MRI protocol. Steps, detailed elsewhere (Jernigan et al. 2016), included a 3D *T*_*1*_-weighted inversion prepared RF-spoiled gradient echo scan using prospective motion correction (PROMO), for cortical and subcortical segmentation; and a 3D *T*_*2*_-weighted variable flip angle fast spin echo scan, also using PROMO, for detection and quantification of white matter lesions and segmentation of CSF.

The CIVET processing pipeline, (http://www.bic.mni.mcgill.ca/ServicesSoftware/CIVET) developed at the Montreal Neurological Institute, was used to compute cortical thickness measurements at 81,924 regions covering the entire cortex. A summary of the steps involved follows; the *T*_*1*-_weighted image is first non-uniformity corrected, and then linearly registered to the Talairach-like MNI152 template (established with the ICBM152 dataset). The non-uniformity correction is then repeated using the template mask. The non-linear registration from the resultant volume to the MNI152 template is then computed, and the transform used to provide priors to segment the image into GM, WM, and cerebrospinal fluid. Inner and outer GM surfaces are then extracted using the Constrained Laplacian-based Automated Segmentation with Proximities (CLASP) algorithm, and cortical thickness is measured in native space using the linked distance between the two surfaces at 81,924 vertices.

Each subject’s cortical thickness map was blurred using a 30-millimeter full width at half maximum surface-based diffusion smoothing kernel to impose a normal distribution on the corticometric data, and to increase the signal to noise ratio.

Quality control (QC) of these data was performed by two independent reviewers. As a result of this process, data with motion artifacts, a low signal to noise ratio, artifacts due to hyperintensities from blood vessels, surface-surface intersections, or poor placement of the grey or white matter (GM and WM) surface for any reason were excluded. In total, 934 unique participants with MRI scans were obtained from PING in a Box. Of these, 905 participants passed quality control procedure. Next, filtering for individuals with information for demographics (age, gender, scanner, SES, ethnicity) and vocabulary scores resulted in a final sample of 757 participants. The demographics of the resulting participants (age, gender, SES, language abilities) used for the study are given in **Table 1**.

**Table 1:** Demographics of the subjects used in the study. Means, with standard deviation given in parentheses. SES = socioeconomic status (measured by parental occupation)

| Scale | Description |
| --- | --- |
| 1 | Unskilled employees |
| 2 | Machine operators and semi-skilled employees |
| 3 | Skilled manual employees |
| 4 | Clerical and sales workers, technicians and owners of little businesses (<2 employees) |
| 5 | Administrative personnel, owners of small businesses and minor professionals |
| 6 | Business managers, proprietors of medium-sized businesses and lesser professionals |
| 7 | Higher executives of large concerns, proprietors and major professionals |

### Statistical Analyses

In order to determine interactive effects of SES (as measured with parental occupation) on cortical thickness with age, general linear models were constructed for each vertex, with the data centered at one-year intervals between three and 21 years. Models with quadratic age terms were found to fit the data significantly better than models with only lower degree age terms, consistent with earlier findings (Noble et al. 2015; Piccolo et al. 2016). Thus, cortical thickness was modelled as:

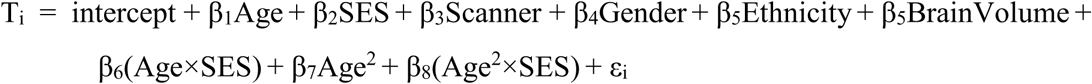

where *i* is a vertex, *Age* is centered as described above, *ε* is the residual error, and the intercept and the *β* terms are the fixed effects. Such a centering approach is similar to earlier published studies investigating developmental trajectories (Shaw et al. 2006, 2007; Nguyen et al. 2013; Khundrakpam et al. 2017). All statistical analyses were done using the SurfStat toolbox (http://www.math.mcgill.ca/keith/surfstat/).

At every cortical point, the *t*-statistic for the association between cortical thickness and SES was mapped onto a standard surface; a random field theory (RFT) correction for multiple comparisons (Worsley et al. 2004) was then applied to the resultant map to determine the regions of cortex showing statistically significant association between cortical thickness and parental occupation. In order to better characterize the age-related patterns of association between cortical thickness and SES, we divided the data sample into two groups: group with Lower SES and group with Higher SES (see **Table 2)** and curves were fitted to the cortical thickness data for the two groups at the peak vertex with the maximum *t*-statistic. Next, we set out to explore age-related differences in cortical thickness for Lower and Higher SES groups. For this, we divided the data sample into 3 age groups: childhood (age = 3 −8.9 years), early adolescence (age = 9 – 13.9 years) and late adolescence (age = 14 −21 years) (see **Table 3**). Note that earlier studies categorized the SES scale 1-3 as the group with Lower SES and SES scale 4-5 as the group with Middle SES (Piccolo et al. 2016; Brito et al. 2017). As can be seen from **Table 3**, the number of participants in Late Adolescence for scale 1-5 is much smaller compared to that of scale 6-7 (86 compared to 200); so, categorizing scale 1-5 to 2 groups would lead to disproportionate number of subjects for group comparisons. In view of this, for our study, we categorized scale 1-5 as the group with lower SES. Considering recent evidence of earlier pubertal milestones particularly in the United States children (Parent et al. 2003; Herman-Giddens 2006), we defined early adolescence as the range of age 9 to 13.9 years. Within each age group, group difference (group with Lower SES vs Higher SES individuals) in cortical thickness was computed for all significant vertices (see preceding section). Within each age group, age, gender, scanner, ethnicity and brain volume were included as covariates and the adjusted cortical thickness was used for the comparative analyses. Since there were nine sites but 12 scanners (with one site with two scanners, and another with three scanners), scanner was put as covariate in the analyses. Since parental occupation was categorical, it was dummy coded in the analyses.

**Table 2:** Details of groups with Lower and Higher SES groups. Means, with standard deviation given in parentheses. SES = socioeconomic status (measured by parental occupation).

|  | **Subjects (N)** | **Age (years)** | **Males/Females** | **SES** |
| --- | --- | --- | --- | --- |
| **Lower SES** | 446 | 3.4 – 21 (11.8 ± 4.9) | 226/220 | 1 – 5 (3.8 ± 0.8) |
| **Higher SES** | 311 | 3.3 – 21 (12.8 ± 4.9) | 163/148 | 6 – 7 (6.4 ± 0.5) |

**Table 3:** Details of groups with Lower and Higher SES for different age groups. Means, with standard deviation given in parentheses. SES = socioeconomic status (measured by family occupation)

|  | **Childhood**<br>**(3 - 8.9 years)** | **Early Adolescence**<br>**(9 - 13.9 years)** | **Late Adolescence**<br>**(14 - 21 years)** |
| --- | --- | --- | --- |
| **Subjects (N)** | Lower SES: 149<br>Higher SES: 80 | 143<br>98 | 86<br>200 |
| **Age (years)** | Lower SES: 6.5 ± 1.7<br>Higher SES: 6.4 ± 1.6 | 11.1 ± 1.5<br>11.5 ± 1.5 | 17.3 ± 2.2<br>17.9 ± 2.1 |
| **Males/Females** | Lower SES: 72/77<br>Higher SES: 45/35 | 75/68<br>52/46 | 45/41<br>101/99 |
| **SES** | Lower SES: 3.7 ± 0.7<br>High-SES: 6.2 ± 0.4 | 3.8 ± 0.8<br>6.5 ± 0.5 | 3.1 ± 0.9<br>5.9 ± 0.8 |
| **Reading scores** | Lower SES: 59.3 ± 41<br>Higher SES: 57.8 ± 43 | 128.3 ± 41<br>137.9 ± 46 | 185.8 ± 50<br>189.9 ± 48 |
| **Vocabulary scores** | Lower SES: -0.5 ± 0.9<br>Higher SES: -0.4 ± 0.9 | 0.7 ± 0.9<br>0.9 ± 1.0 | 1.9 ± 1.0<br>1.9 ± 0.9 |

A similar analysis was performed for each age group comparing the group difference (group with Lower SES vs Higher SES individuals) in language abilities (vocabulary and reading scores). Within each age group, age, gender, site, ethnicity and brain volume were included as covariates and the adjusted scores were used for the comparative analyses.

Next, based on previous findings (Noble et al. 2015; Farah 2017), we investigated whether the link between SES and cognition (language abilities) is mediated by cortical thickness (at the peak vertex). Since we set out to explore non-linear/quadratic effect of age on association between SES and cortical thickness, we extended the mediation analysis to include ‘moderation’ by age and age^2^. First introduced by James and Brett (James and Brett 1984), a moderator effect arises when the effect of one predictor is changed by a second predictor meaning an interaction effect emerges. In our case, the association between SES and language abilities is mediated by brain structure; this mediation in turn may vary as a function of age and age^2^. To fully disentangle the nature of the relationships between the variables, it is necessary to combine these two approaches – mediation and moderation. Moderated mediation analysis was performed according to Hayes (Hayes 2015), by implementing the regression formulae in R, using ‘lavaan’ and ‘processr’ packages.

## Results

### Non-linear (SES × age) interaction with cortical thickness

Significant positive association (*p* < 0.05, RFT-corrected) between SES and cortical thickness was observed during the age period 9 −13 years in several brain regions located in the left frontal, temporal, parietal and occipital cortices, and the right parietal cortex (**Figure 1**). There was no significant negative association of SES and cortical thickness during any of the other age periods.

**Figure 1:**
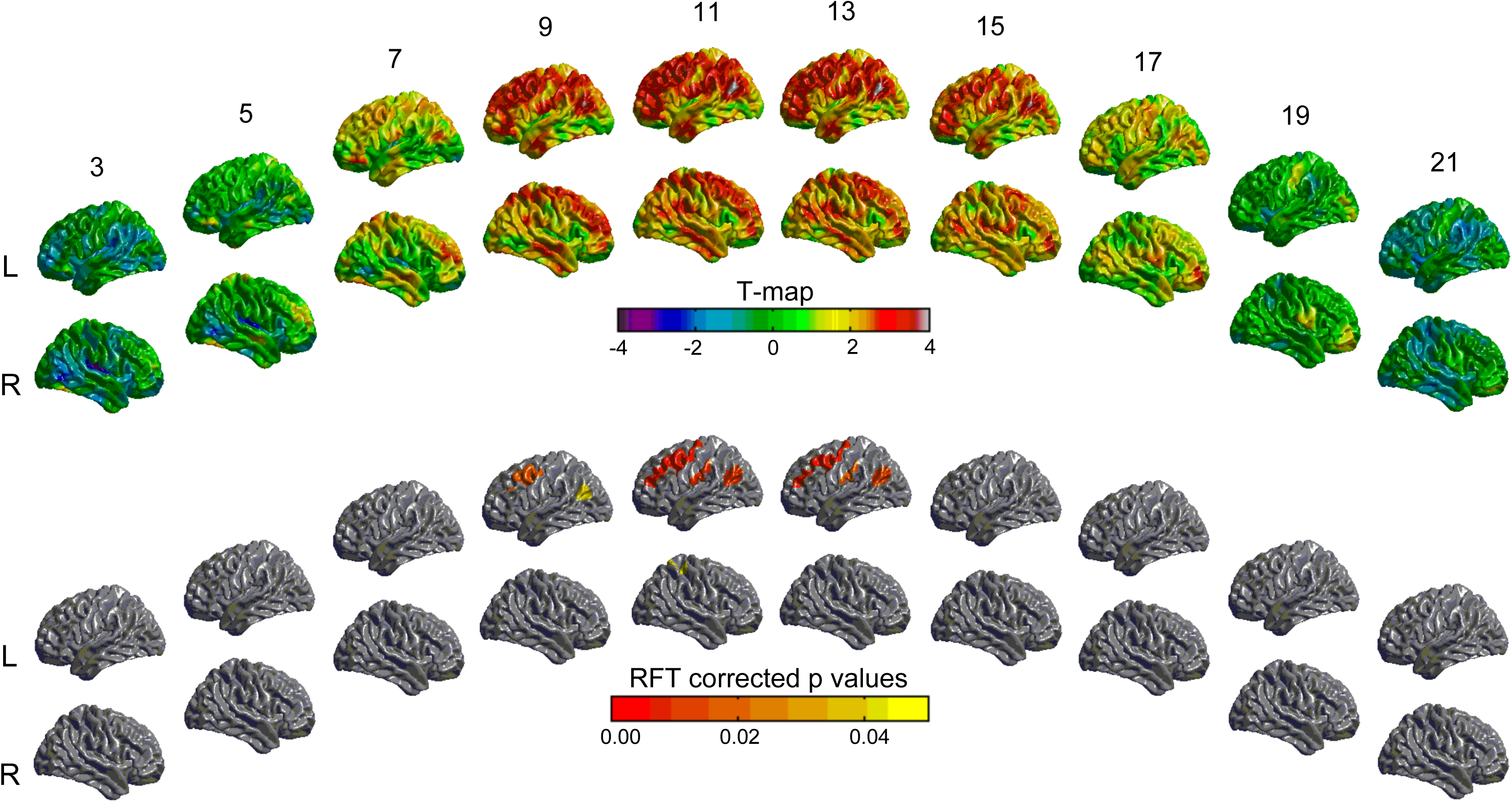
Interactive effects of age and socioeconomic status (SES) on cortical thickness. The *t-*statistics (across vertices on the surface) and multiple comparison-corrected *p-*statistics (across age bins) for the association of cortical thickness and SES (measured by family occupation) across time. The statistics are shown as surface maps, with lateral views of the left and right hemispheres shown for both statistics. The upper two rows of surface maps show the *t-*statistics for the association at ages ranging from 3 to 21 years of age, with age increasing from left to right, and the left hemisphere shown above the right hemisphere. The lower two rows show the significant *p-*statistics (*p* < 0.05, RFT-corrected for multiple comparisons across the age bins, see **Methods**) for the association. Note that there is significant positive association between cortical thickness and family occupation during age 9 to 13 years in regions of the left frontal, temporal, parietal and occipital cortex, and the right parietal cortex. No significant associations are observed thereafter. *L* and *R* denote left and right hemisphere, respectively. The numbers above the top row of surface maps indicate the age (in years) for the statistics depicted in that column.

### Age-related difference in cortical thickness for group with Higher compared to lower SES

In order to better characterize the non-linear (SES × age) interaction with cortical thickness, we plotted cortical thickness at the peak vertex (*T* = 4.45, MNI coordinates: x = −55, y = −64, z = 25) for the groups with Lower and Higher SES individuals (**Figure 2**). Qualitatively, the largest dissociation between the fitted curves for two groups was observed around 13 years, and the curves merged around 19 years.

**Figure 2:**
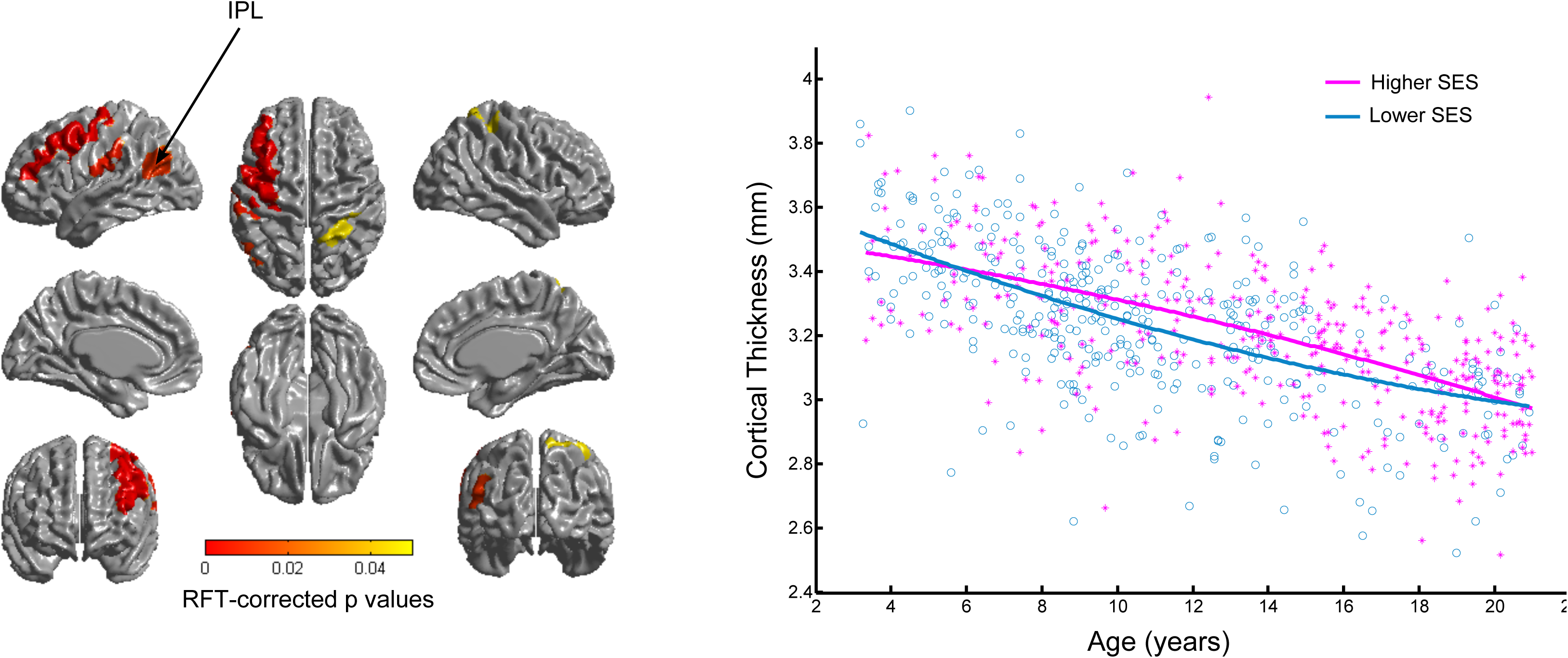
Fitted curves of cortical thickness in groups with Lower and Higher SES. The centre panel shows a surface map of the cortical regions for which there were significant group differences in cortical thickness (group with Higher and Lower SES). The scatter plots around the periphery show the curves fit to the cortical thickness data for each group at the peak vertex with the maximum *t*-statistics (MNI coordinates: x = −55, y = −64, z = 25, see **Methods**). Note the largest dissociation between the fitted curves for groups with Lower and Higher SES was observed around 13 years, and the curves merged around 19 years. Note, x-axis = age (years), y-axis = cortical thickness (mm), SES = socio-economic status.

### SES-related group difference in cortical thickness for age groups

Significantly greater cortical thickness was observed for group with higher SES compared to Lower SES during childhood (*T* = 2.63, *df* = 227, *p* = 0.009) and early adolescence (*T* = 6.57, *df* = 239, *p* = 0). Note, the extent of group difference in cortical thickness was much larger during early adolescence as compared to childhood (**Figure 3**).

**Figure 3:**
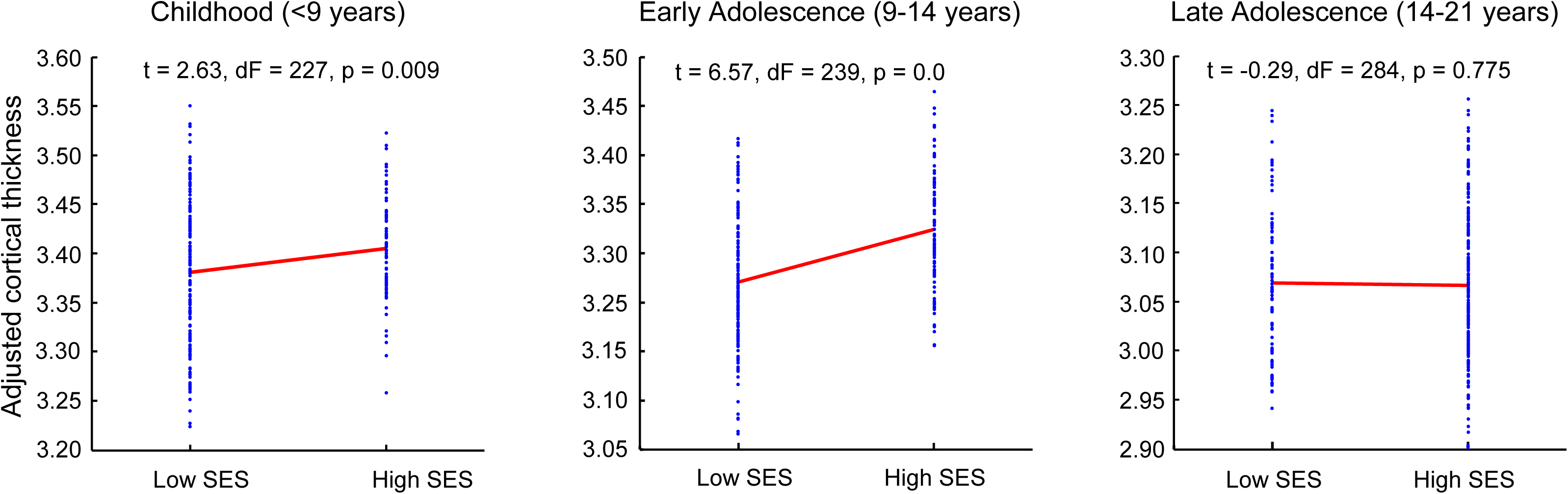
Group difference (Lower vs Higher SES) in cortical thickness across age groups. Group comparison in cortical thickness for groups with Higher and Lower SES for the age groups – childhood (3-8.9 years), early adolescence (9-13.9 years) and late adolescence (14-21 years). Note significantly greater cortical thickness in Higher compared to Lower SES during early adolescence (to a great extent), and during childhood (lesser extent). There was no significant group difference in cortical thickness during late adolescence. Note, x-axis = age groups, y-axis = adjusted cortical thickness, SES = socio-economic status.

### SES-related group difference in language abilities for age groups

Significantly greater vocabulary (*T* = 4.81, *df* = 239, *p* = 0) and reading scores (*T* = 4.14, *df* = 239, *p* = 0) were observed for group with Higher SES compared to Lower SES during early adolescence. There was no significant group difference for both the scores during childhood and late adolescence (**Figure 4**).

**Figure 4:**
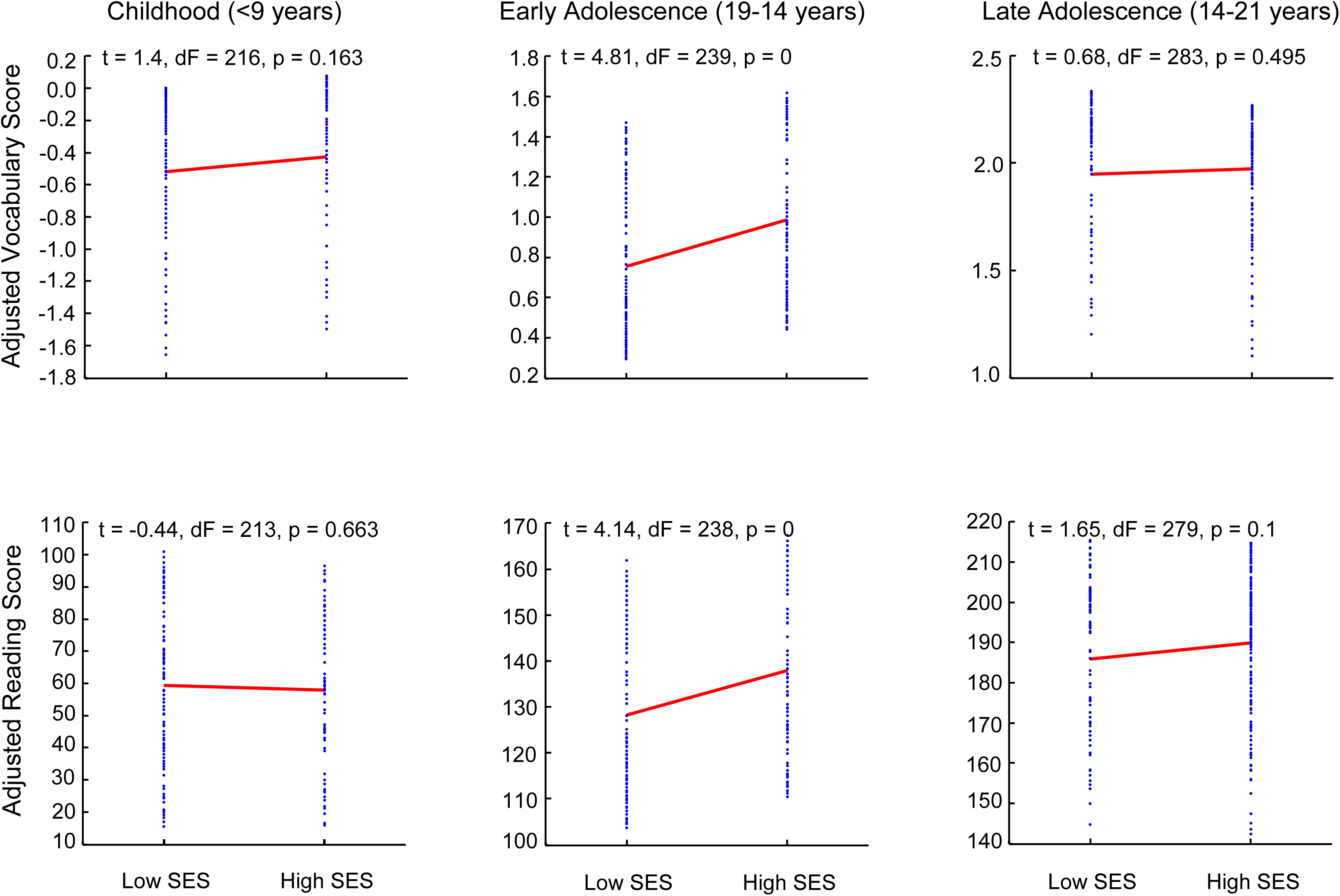
Group difference (Lower vs Higher SES) in language abilities across age groups. Group comparison in language abilities (vocabulary and reading scores) for groups with Lower and Higher SES for the age groups – childhood (3-8.9 years), early adolescence (9-13.9 years) and late adolescence (14-21 years). Note, significantly greater vocabulary (upper row) and reading (lower row) scores for groups with Higher compared to Lower SES during early adolescence. There was no significant group difference for both the scores during childhood and late adolescence. Note, x-axis = groups, y-axis (upper row) = adjusted vocabulary score, y-axis (upper row) = adjusted reading score, SES = socio-economic status.

### Age nonlinearly moderates mediation of SES, cortical thickness and language abilities

Using a mediation model, we first investigated the extent to which cortical thickness (at peak vertex) mediated the link between SES and language abilities (vocabulary scores). The direct effect of SES on vocabulary was *β* = 0.03, *p* < 0.003, indicating a weak but significant association between SES and vocabulary scores (**Figure 5**). This effect was reduced to *β* = 0.01, *p* < 0.003 when controlling for cortical thickness. The direct effect of SES on cortical thickness was *β* = 0.06, *p* < 0.003, while that of cortical thickness on vocabulary scores was *β* = 0.05, *p* < 0.003. Next, we explored whether *age* and *age*^*2*^ moderated the links between SES, cortical thickness and vocabulary scores. The effect of *age* on SES-cortical thickness link was *β* = 0.37, *p* < 0.003, while that of *age*^*2*^ was *β* = −0.40, *p* < 0.003 indicating that association of SES and cortical thickness linearly increased with age, but was countered by a negative effect of *age*^*2*^, resulting to a quadratic trajectory with greatest SES-cortical thickness association around the middle of the age range of the data sample, which largely corresponded to early adolescence (9-13.9 years).

**Figure 5:**
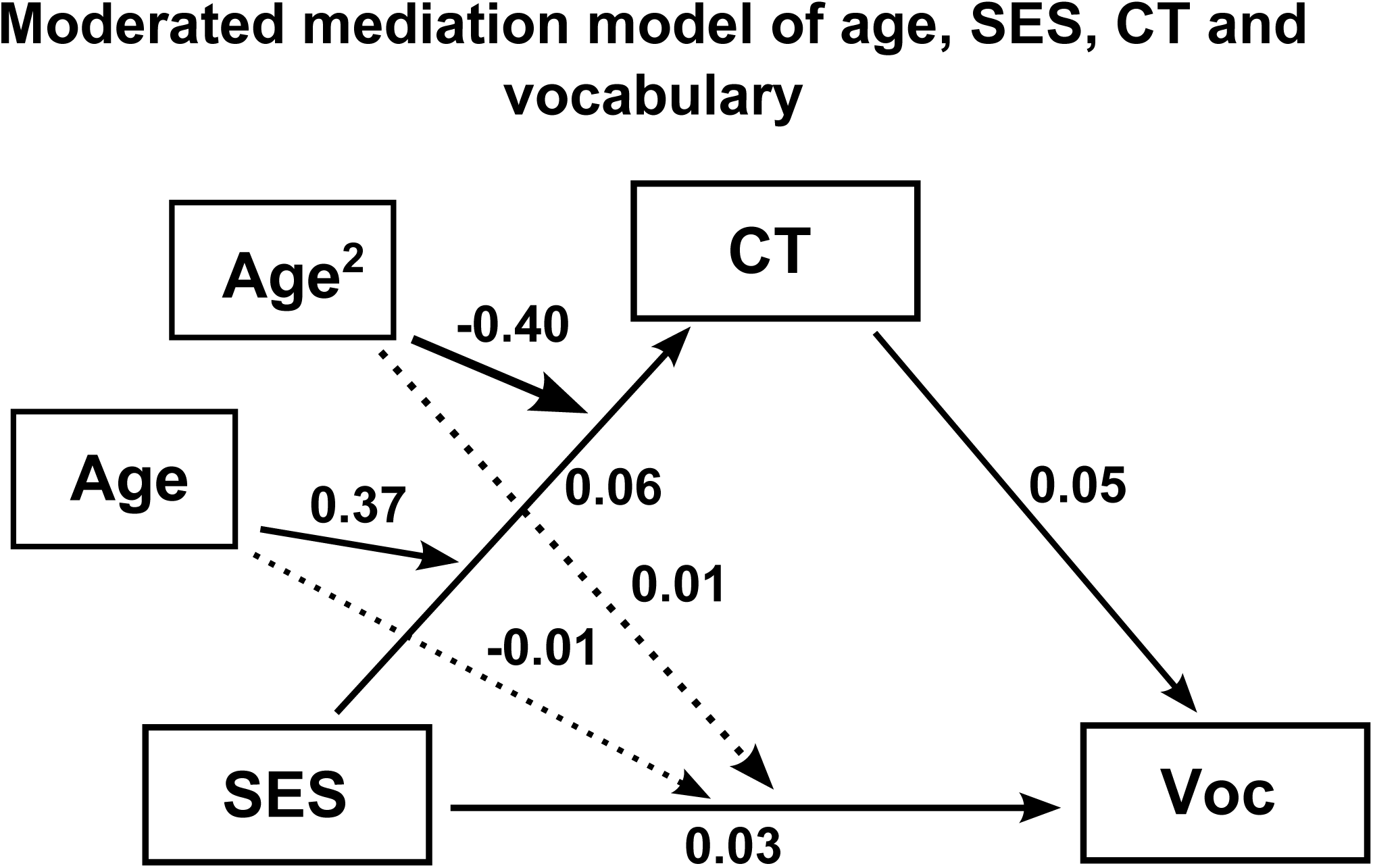
Age nonlinearly moderates mediation of SES, cortical thickness and language abilities: Cortical thickness mediated the link between SES and language abilities (vocabulary scores). This mediation was in turn moderated by *age* and *age*^*2*^ (see **Results**) in such a way that association of SES and cortical thickness linearly increased with age, but was countered by a negative effect of *age*^*2*^, resulting to a quadratic trajectory with greatest SES-cortical thickness association around the middle of the age range of the data sample, which largely corresponded to early adolescence (9-13.9 years). Note, CT = cortical thickness, Voc = vocabulary scores, SES = socio-economic status.

## Discussion

In this study, using data from a large sample of typically-developing children and adolescents with ages ranging from 3 to 21 years, we showed non-linear (SES × age) interaction with cortical thickness. Specifically, we observed a significant positive association between SES and cortical thickness during the age period of 9-13 years in several regions including the left frontal, temporal, parietal and occipital cortices, and the right parietal cortex. The nonlinear interaction of age and SES was better illustrated by splitting the data into two groups (subjects from Lower and Higher SES families), and estimating age-specific group differences in cortical thickness maps by centering the data at one-year intervals between 3 and 21years. We observed significantly greater cortical thickness for subjects from Higher SES families during early adolescence, but not during childhood or late adolescence. Interestingly, significantly greater language abilities (assessed with reading and vocabulary scores) were observed for the group with Higher SES individuals during early adolescence, suggesting a link between SES, cortical thickness and language abilities. Indeed, we observed that cortical thickness mediated the link between SES and language abilities, and this mediation was moderated by age/age^2^ in a quadratic pattern, indicating a larger effect during early adolescence. Our results, drawn from cross-sectional data, provide a basis for further longitudinal studies to test the hypothesis that early adolescence may be a sensitive time window for the impact of SES on brain and neurocognitive development.

Consistent with our data, recent studies have indicated that the impacts of SES on the brain changes with age. In a longitudinal study on infants and toddlers (aged 5 months to 4 years), children from low-SES showed slower trajectories of cortical growth compared to that of high-SES during infancy and early childhood (Hanson et al. 2013). Another study using the PING dataset reported an interaction of SES and age such that higher SES was associated with greater volume in the left superior temporal gyrus and left inferior frontal gyrus of participants during adolescence (Noble et al. 2012). More recently, in another study using a large data sample, the same group observed nonlinear (curvilinear) cortical trajectories for participants with lower SES and linear trajectories for participants with higher SES (Piccolo et al. 2016). However, this *non-linear* (SES × age) interaction was observed for the average cortical thickness (of all brain regions) whereas at region-level, this non-linear interaction was not significant. As distinct from those data, in our study, we observed significant non-linear (SES × age) interaction with cortical thickness at several brain regions located in the left frontal, temporal, parietal and occipital cortices, and the right parietal cortex (**Figure 1**). Additionally in our study, at the peak vertex with maximum *T*-score for the interaction, SES was associated with language abilities, and this association was in turn mediated by cortical thickness (**Figure 5**). Taken together, our study (using parental occupation as a proxy for SES) extends previous findings of non-linear (SES × age) interaction with cortical thickness by showing high regional specificity and by showing a link of this interaction with cognition. It may be noted that the differences of our findings from that of previous studies could result from different measures of SES (parental education, family income vs parental occupation), and different MRI preprocessing pipelines. Indeed, supplementary analysis using parental education (**Supplementary Figure 1**) and family income (**Supplementary Figure 1**) largely replicated findings from (Piccolo et al. 2016) reiterating that the findings reported in the study may be unique for parental occupation.

Interestingly, our findings of cortical regions showing significant interactions of SES and age were asymmetric, predominantly localized in left hemisphere, including language-related regions around the arcuate fasciculus (**Figure 3**). In fact, the peak vertex with maximum *T*-score for the interaction was localized at the inferior parietal cortex, part of the posterior speech area. Coupled to this, there was direct association of SES and language abilities mediated by cortical thickness. Our findings further add to the accumulating evidence from functional and structural imaging studies in support of the purported role of the brain in mediating the effects of SES on certain life outcomes (Noble et al. 2005, 2006, 2012; Farah 2017). In particular, these studies point to key effects on language development and the underlying neural circuitry. Electroencephalographic (EEG) studies have found evidence of a maturational lag in the prefrontal cortex (Otero 1997) in addition to left-frontal hypoactivity (Tomarken et al. 2004) in lower SES preschool children and adolescents, respectively; these findings correspond to the findings of behavioral studies examining language and attention differences between groups (Hackman and Farah 2009). A review of fMRI studies suggest decreased functional specialization in language regions for low-SES kindergartners (Raizada et al. 2008), less mature frontal gamma power in low-SES children (Tomalski et al. 2013), and decreased functional connectivity at resting-state in low-SES children and adults (Sripada et al. 2014; Barch et al. 2016; Marshall et al. 2018), suggesting a comparative delay in functional brain development (Tooley et al. 2018). In agreement with these functional studies, our findings of asymmetric cortical correlates of SES disparities, with reduced thickness for participants with lower SES in the left-hemispheric language-related regions with concurrent decrease in vocabulary and reading scores also suggest delayed cortical development with behavioural consequences for participants with lower SES.

In terms of understanding the underlying mechanisms behind the distinct trajectories for groups with Lower and Higher SES individuals, we can leverage the knowledge of neurodevelopmental trajectories that have been useful in understanding normal and abnormal brain development (Giedd et al. 1999; Shaw et al. 2006, 2007, 2008; Gogtay et al. 2008; Zielinski et al. 2014; Khundrakpam et al. 2017). In light of these studies, we can interpret the neurodevelopmental trajectories for group with Lower SES individuals as deviant trajectories with faster thinning during childhood and leveling off in adolescence, corresponding with earlier findings (Piccolo et al. 2016). These results align with studies using animal models that have demonstrated mechanisms of early adversity and deviant brain trajectories (Bath et al. 2016; Fareri and Tottenham 2016). Specifically, early adversity has been associated with processes such as increased cell death, altered neuronal morphology (Bath et al. 2016; Callaghan and Tottenham 2016), which in turn may be reflected in the faster cortical thinning during childhood for the group of individuals from Lower SES families.

Our findings indicate that the impact of SES on brain structure and cognitive development may be most salient during early adolescence. Although speculative and pending future longitudinal studies, our findings may align with evidence from developmental cognitive neuroscience showing reorganization of brain structure in terms of synaptic pruning and myelination processes with concomitant changes in brain function that have led to an increasingly convincing view of adolescence as a window of major neurocognitive plasticity. These data, consistent with evolutionary life history accounts of development, suggest that the adolescent brain is especially susceptible to social environmental stimuli, leading many to posit that this age period is a critical period for the biological embedding of socio-cultural input. MRI findings point to an inverted-U shaped pattern of grey matter development in multiple regions of the brain including frontal, temporal and parietal cortex (Giedd et al. 1999; Gogtay et al. 2004; Shaw et al. 2008), such that grey matter volume peaks around the age of puberty onset, before a process of synaptic pruning begins into mid-adolescence and early adulthood. Although evidence of the precise mechanisms remains tentative, animal studies and neuroimaging research suggests that there is an interaction between pubertal hormones and structural brain development (Sisk and Zehr 2005; Cahill 2006; Herting and Sowell 2017). Puberty, which marks the beginning of adolescence and is characterized by dramatic changes in hormones, emotions, physical growth and social life, is a particularly important moment for the brain’s contextual sensitivity (Sisk and Foster 2004; Blakemore et al. 2010). For example, gonadarche, which is initiated between ages 8 and 14 years in females, and between ages 9 and 15 in males, begins with the activation of the hypothalamic-pituitary-gonadal (HPG) axis (Abreu and Kaiser 2016). The mechanisms that activate the HPG axis are sensitive to factors such as the quality of available nutrition and the presence of pathogens (Worthman and Trang 2018). Stressed ecologies during early life, in the form of low socioeconomic position or poor parenting styles, have been shown to impact on the timing and events of the pubertal process, such that stressful early life events cue individuals both physiologically and behaviorally to begin investment in reproduction sooner (Belsky and Shalev 2016). Further longitudinal studies incorporating pubertal hormone measures from participants during childhood through adolescence could shed light on the possibility the current data raise that like the window during the early years, the brain may also be sensitive to contextual adversity during the peri-pubertal age period. Building on developmental psychology and anthropology, longitudinal data from large cohorts of young people could determine the role of the brain in regulating the impacts of poverty and subsequent trajectories of brain maturational processes and cognitive development.

In our study, we specifically used parental occupation as a proxy for social standing. Parental occupation is one of the three most commonly used proxies for SES, along with family income and parental education. It is important to note that there is great debate around the inconsistency in SES measures. The lack of consistency raises questions about the degree to which these studies (using either a single proxy or composite, multivariate representation of poverty) can be accurately synthesized or compared. The three components of SES are statistically correlated and conceptually related in complex ways (Braveman et al. 2005). For instance, a successful ballet dancer may have low educational attainment (in terms of number of years or higher degrees) but high occupational prestige, or, a professor may have more education and occupational prestige than a mechanic, but lower income. Composite measures of SES such as the frequently used Hollingshead four-factor index, while commonly used, may obscure distinct processes since the constituent factors (income, occupation, education) correspond to different lived experiences and neural outcomes. Household income level is most commonly used in the literature but defining a child’s SES solely from a material standpoint obscures proximal factors that may be better predictive factors of brain impingement such as exposure to environmental toxins or maternal stress, both of which have demonstrated impacts on child cognitive development and yet are not reflected in a definition weighted to purchasing power (D’Angiulli et al. 2012). Furthermore, income information does not capture the fact that people (especially low-income groups) may have income in kind, such as food stamps, or crops which are traded. Income can also be an unreliable indicator of social standing for self-or transitorily employed workers (McKenzie 2005). While parents’ number of years in education has consistently shown relationships with cognitive outcomes, it may mask quality of education or the resultant occupational prestige as illustrated above. In view of these caveats, and in light of the fact that much of the neuroimaging literature has focused only on parental income and education, we propose that parental occupation may also be a sensitive indicator of childhood and adolescent SES because it captures position in the social hierarchy, which has consistently been shown to be intimately related to health and life chances (Marmot et al. 1978, 1991; Pinilla et al. 2017).

Our findings of null results (no difference for groups with lower and higher SES) for language abilities in late adolescence may seem counterintuitive given recent reports of increased language abilities for subjects with higher SES across childhood and adolescence (Brito et al. 2017). As elaborated in **Supplementary Text**, a closer investigation of the association of SES, cortical thickness and language abilities revealed that participants with extremes of cortical thickness (greatest and lowest cortical thickness) show minimal SES-related group difference in language abilities. This in turn suggests (considering cortical thickness negatively correlates with age) that participants with extreme ages (youngest and oldest) would show minimal SES-related group difference in language abilities. In other words, it is possible that at the group-level, the language abilities may not differ for individuals with lower and higher SES during early childhood and late adolescence (for details, see **Supplementary Text**).

The main limitation of our study is the use of cross-sectional data; as such, our findings are correlational rather than causal, and must therefore be interpreted cautiously. It is therefore not clear whether SES disparities lead to lesser language abilities via altered neurodevelopmental trajectories. Nonetheless, the analysis of a large cohort of young people is suggestive of differential effects of SES during development. Of course, further investigations are required to distinguish between differential sensitivity and differential rates of maturation during adolescence, and to unpack the underlying biological mechanisms and social factors such as family stress, prenatal factors, cognitive deprivation or toxins. Future studies utilizing longitudinal MRI data from available datasets such as the ABCD (Casey et al. 2018), tracking children over time with social and environmental measures (Zucker et al. 2018), along with animal studies of the possible underlying biological pathways will help better understand the complex interplay of SES, brain and life outcomes (Hackman et al. 2010).

Using a large sample of typically developing children and adolescence, our study demonstrated a nonlinear (quadratic) interaction of age and SES (measured by parental occupation) on cortical thickness, such that individuals from Higher SES families were associated with having greater cortical thickness (predominantly in left-hemispheric cortical regions including language-related areas) as well as greater language abilities during early adolescence. Our findings, drawn from cross-sectional data, provide a basis for further longitudinal studies to test whether early adolescence is a sensitive period for social contextual factors on brain development and cognitive outcomes.

