## Supplementary material for "Non-linear effects of socioeconomic status on brain development: associations between parental occupation, cortical thickness and language skills in childhood and adolescence": Supplemenatry Material

### for

### ‘Socioeconomic status moderates age-related differences in cortical thickness and language abilities during childhood and adolescence’

**Supplementary Figure S1:**

**Interactive effects of age and parental education on cortical thickness.** The *t*-statistics (across vertices on the surface) for the association of cortical thickness and SES (measured by parental education) across time. The statistics are shown as surface maps, with lateral views of the left and right hemispheres shown for both statistics. The upper two rows of surface maps show the *t*-statistics for the association at ages ranging from 3 to 19 years of age, with age increasing from left to right, and the left hemisphere shown above the right hemisphere. The lower two rows show the significant *p*-statistics ($p < 0.05$, RFT-corrected for multiple comparisons across the age bins) for the association. Note that there is significant positive association between cortical thickness and family occupation during age 11 to 13 years in regions of the right temporal cortex. No significant associations are observed thereafter. The numbers above the top row of surface maps indicate the age (in years) for the statistics depicted in that column.

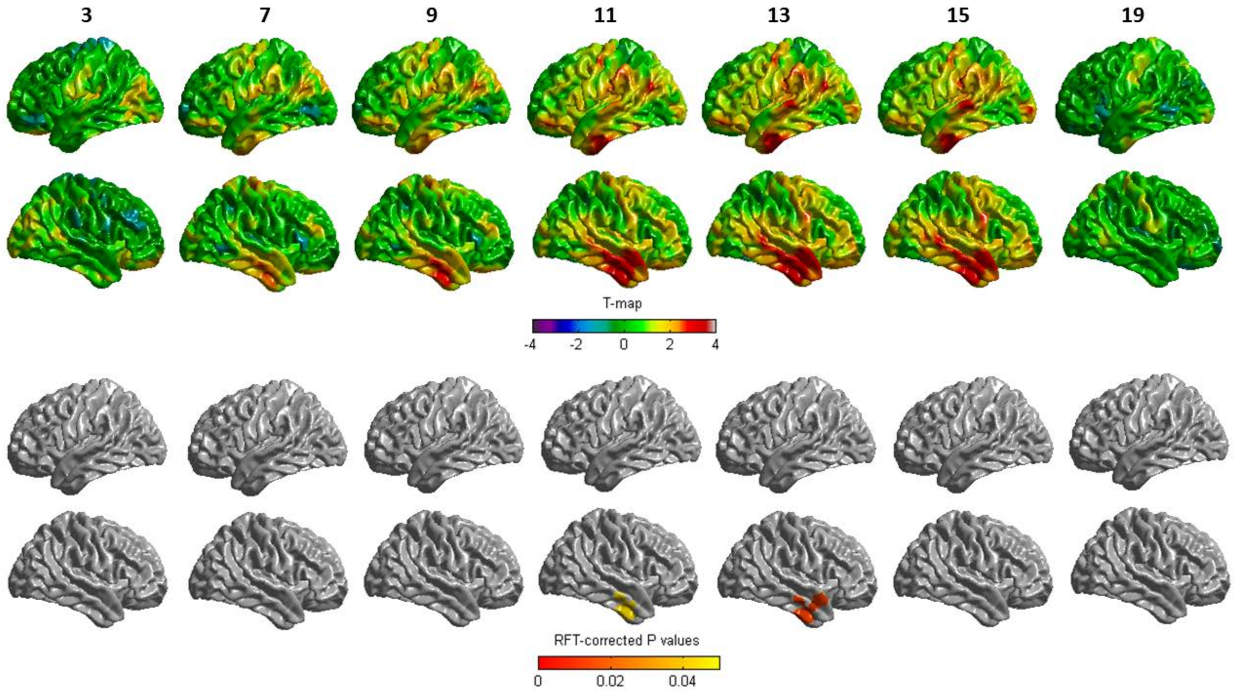

**Supplementary Figure S2:**

**Interactive effects of age and family income on cortical thickness.** The *t*-statistics (across vertices on the surface) for the association of cortical thickness and SES (measured by family income) across time. The statistics are shown as surface maps, with lateral views of the left and right hemispheres. The row of surface maps show the *t*-statistics for the association at ages ranging from 3 to 19 years of age, with age increasing from left to right, and the left hemisphere shown above the right hemisphere. The numbers above the top row of surface maps indicate the age (in years) for the statistics depicted in that column. Note that there is no significant association at any time point.

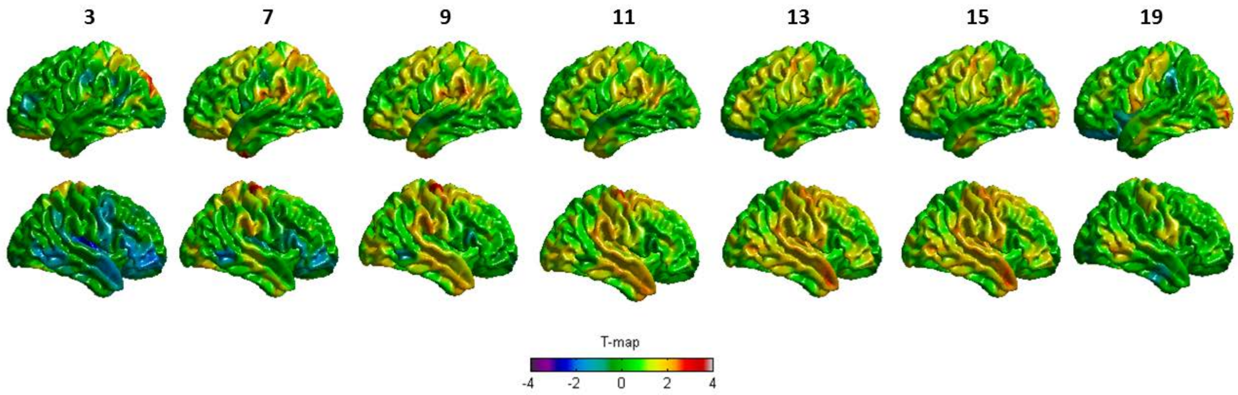

**Supplementary Text:**

Findings from Brito et al. 2017 revealed association of SES, cortical thickness and language abilities.

Association of family income, cortical thickness and reading:

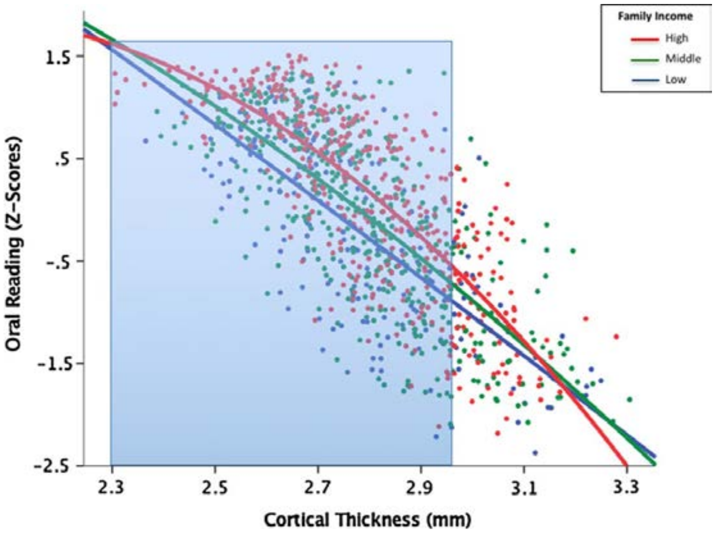


Association of family education, cortical thickness and vocabulary, reading:

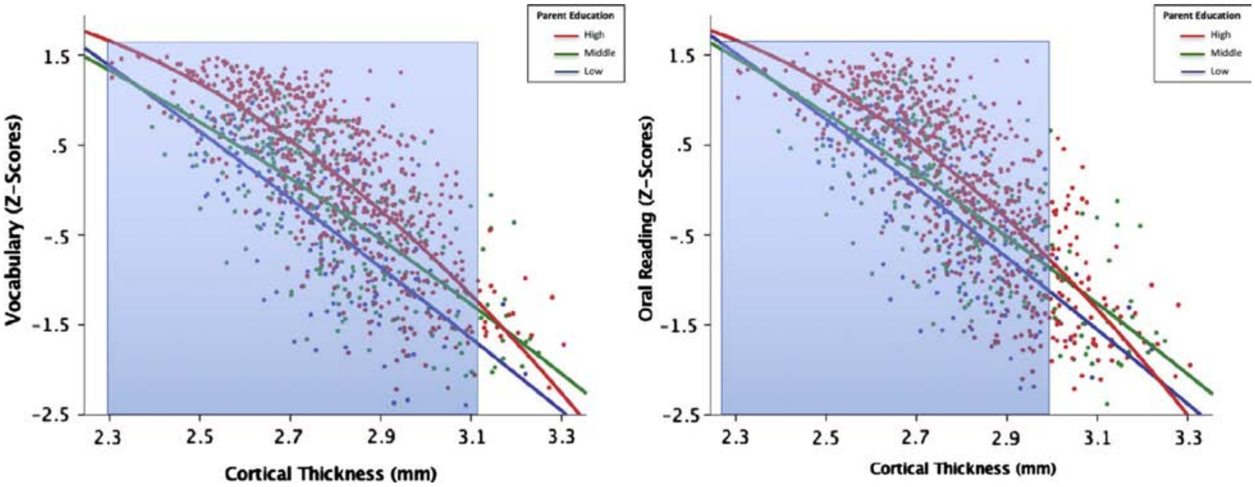

In our study, we also observed similar association of SES, cortical thickness and language abilities.

Association of family occupation, cortical thickness and vocabulary, reading:

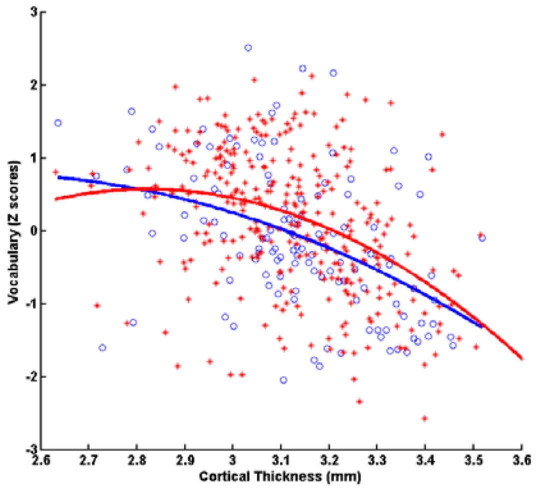


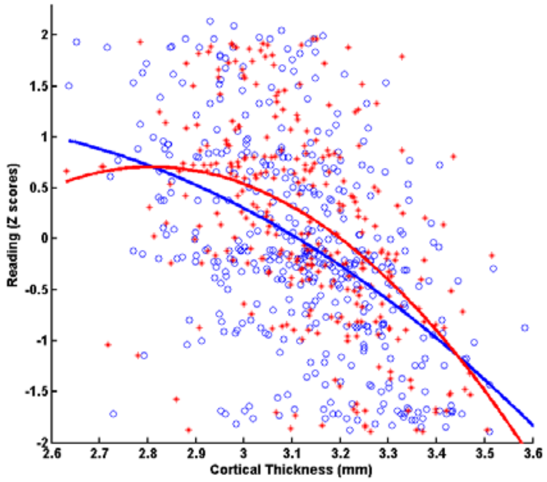


Cortical thickness negatively correlates with age.

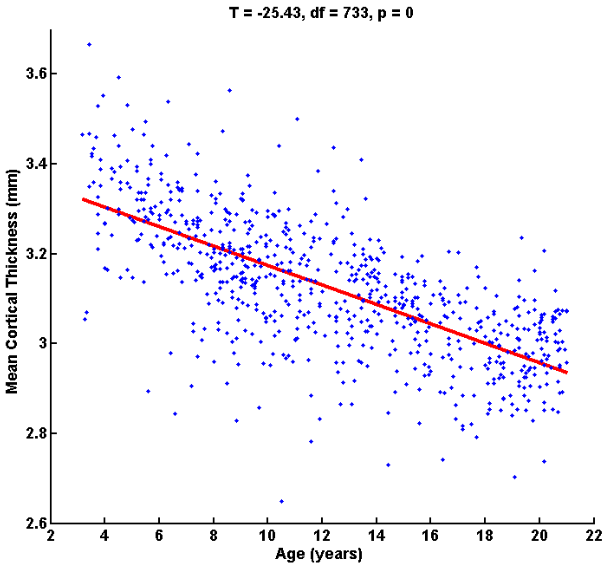


Thus, in general, subjects with greater cortical thickness are younger children and subjects with lesser cortical thickness are older children (adolescents). The previous observations (both ours and that of Brito et al.) may reflect the following association –

Representative association of SES, age and language abilities

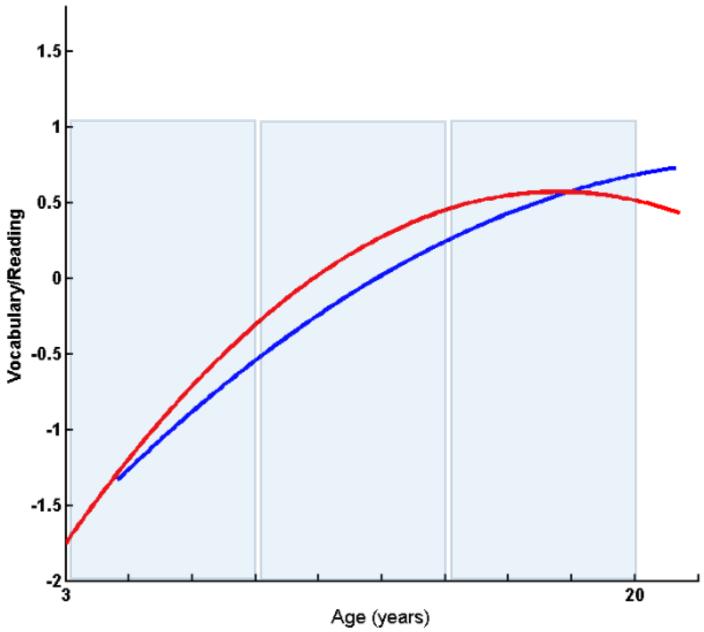


**Childhood Early Adolescence Late Adolescence**

In such a scenario, group difference (subjects with lower and higher SES) in language abilities will be most likely observed for early adolescence. In late adolescence, any initial increased language abilities for higher SES would be offset by decreased abilities in later ages, thereby producing a null group difference.
